# Sex differences in early and term placenta are conserved in adult tissues

**DOI:** 10.1101/2022.08.08.503197

**Authors:** Kimberly C. Olney, Seema B. Plaisier, Tanya N. Phung, Michelle Silasi, Lauren Perley, Jane O’Bryan, Lucia Ramirez, Harvey J. Kliman, Melissa A. Wilson

## Abstract

**Background:** Pregnancy complications vary based on the fetus’s genetic sex, which may, in part, be modulated by the placenta. Further, developmental differences early in life can have lifelong health outcomes. Yet, sex differences in gene expression within the placenta at different time points throughout pregnancy and comparisons to adult tissues remains poorly characterized.

**Methods:** Here, we collect and characterize sex differences in gene expression in term placentas ( ≥ 36.6 weeks; 23 male XY and 27 female XX). These are compared with sex differences in previously collected first trimester placenta samples and 42 non-reproductive adult tissues from GTEx.

**Results:** We identify 268 and 53 sex differentially expressed genes in the uncomplicated late first trimester and term placentas, respectively. Of the 53 sex differentially expressed genes observed in the term placentas, 31 are also sex differentially expressed genes in the late first trimester placentas. Furthermore, sex differences in gene expression in term placentas are highly correlated with sex differences in the late first trimester placentas. We found that sex differential gene expression in the term placenta is significantly correlated with sex differences in gene expression in 42 non-reproductive adult tissues (correlation coefficient ranged from 0.892 to 0.957), with the highest correlation in brain tissues. Sex differences in gene expression were largely driven by gene expression on the sex chromosomes. We further show that some gametologous genes (genes with functional copies on X and Y) will have different inferred sex differences if the X-linked gene expression in females is compared to the sum of the X-linked and Y-linked gene expression in males.

**Conclusions:** We find that sex differences in gene expression are conserved in late first trimester and term placentas and that these sex differences are conserved in adult tissues. We demonstrate that there are sex differences associated with innate immune response in late first trimester placentas but there is no significant difference in gene expression of innate immune genes between sexes in healthy full term placentas. Finally, sex differences are predominantly driven by expression from sex-linked genes.

**Highlights:**

- Sex differences in gene expression in late first trimester placentas are positively correlated with sex differences in gene expression in full term placentas; sex differences develop early and are maintained.
- Sex differences in gene expression on the sex chromosomes in the placenta are correlated to sex differences in adult tissues.
- Sex-linked gametolog genes require additional methodological approaches for accurate quantification.

## Background

The placenta is a specialized organ that is part of and supports a developing fetus by facilitating nutrient transfer, waste removal, and immune regulation with the maternal body (Burton et al, 2015). Trophoblast cells are the first cells to differentiate from the fertilized oocyte forming the outer layer of the blastocyst and invade into maternal uterine tissue forming an interface that grows with the developing embryo and fetus throughout pregnancy. Signals from the placenta elicit proinflammatory TH1 responses in the first and third trimester to create an environment free from pathogens for implantation and prepare the fetus for interaction with the outside world, while an anti-inflammatory response occurs in the second trimester to prevent rejection of the fetus as it undergoes rapid growth and development (Mor et al., 2011). Changes in placental development can have major effects on the fetus and many studies have demonstrated that altered placenta function is associated with pregnancy complications such as preeclampsia and intrauterine growth restriction (Haram et al., 2020). Healthy placental tissue may provide a common reference for studying the developmental origins of sex differences in adverse health outcomes that arise during pregnancy that have continued effects after birth (Burton et al., 2016; Goldstein et al., 2020; Keleher et al., 2021).

Sex-specific differences have been observed in the placenta, the developing fetus, infants, and adults. The specific placental biomarkers s-FLt1, PAI-2, and PLGF have been measured at higher concentrations in female placenta compared to male placenta (Brown et al., 2014), while male placentas have been reported to have higher expression of induced TNF-a response involved in inflammation relative to female placentas (Yeganegi et al., 2009). Male fetuses have a larger crown-to-rump length than females in the first trimester and a slower growth rate of head circumference, while male neonates are heavier than female neonates at birth (Brown et al., 2014). Further, a recent study of adult tissues from the Genotype-Tissue Expression Project (GTEx) demonstrated that sex influences gene expression and cellular composition throughout the body with 37% of all genes showing sex differences in expression in at least one tissue (Oliva et al., 2020). In mouse, chromosomal sex contributes, along with organizational gonadal sex and activational hormones, to sex differences in gene expression in adult tissues (Blencowe et al., 2022) in adult tissues. However, sex differences in early formed placental tissues have not been compared with sex differences in adult tissues.

Therefore, we studied sex differences in the placental gene expression in early and late stage pregnancy and compared with adult tissues. We generated RNA and DNA from 30 male and 30 female term ( ≥ 36.6 weeks) placentas from uncomplicated births. We compared sex differences in term uncomplicated placentas to late first trimester placenta samples (Gonzalez et al., 2018) and to 42 non-reproductive adult tissues to better understand the consistency of sex differential expression across the life span. We found that sex differences in gene expression in term placentas are highly correlated with sex differences in the late first trimester placentas. We found that genes related to immune function are sex differentially expressed in the late first trimester but not at full term. Genes on the sex chromosomes, but not the autosomes, tend to show similar sex differences between the placenta and adult tissues, with the highest correlations between the placenta and brain tissues. Furthermore, we discuss approaches for quantifying gametologous gene expression (expression from genes with functional copies on the X and Y chromosomes due to shared ancestry). Understanding sex-specific changes in placental gene expression during and after pregnancy can provide insight on the mechanisms underlying human growth and development *in utero* and beyond in health and disease.

## Methods

### Samples

Working with the Yale University Reproductive Sciences (YURS) Biobank, we collected 60 term ( ≥ 36.6 weeks) placentas from uncomplicated pregnancies, 30 assigned female at birth and 30 assigned male at birth. Individuals were chosen for inclusion in this study based on self-reported race – Black (33%), White (33%), Asian (23%), and Hispanic (7%) – to increase the likelihood of a genetically diverse sample (Additional Table 1; see demographic analysis below). The placenta samples here were carefully selected to represent the fetal component of the placenta. Tissue samples were taken midway between the chorionic and basal plates from the periphery of the lobules avoiding maternal tissue. Placentas were oriented maternal side up and the basal plate was removed to remove the maternal tissue. Four tissue samples weighing roughly 50 mg were placed in a cryovial, snap frozen in liquid nitrogen, and stored at −80°C until processed for sequencing. Chosen sampling sites were free of visible infarction, calcification, hematoma, and tears. Three samples were obtained from each placenta, one for whole exome sequencing, and two tissue samples from opposing quadrants for RNA sequencing (RNAseq) for a total of 120 RNAseq placenta samples. The placentas were collected and sequenced at two different times, with 12 male and 12 female placentas in the first batch and 18 male and 18 female placentas in the second batch. All placenta samples were collected immediately following live birth via cesarean section (CS) except for one male placenta, which was collected following spontaneous vaginal delivery (SVD), sample ID YPOPS0007M. However, the spontaneous vaginal delivery sample was removed from the study due to failed GC content (Additional Table 1).

### RNAseq data processing

RNAseq libraries were constructed using Illumina TruSeq reverse forward stranded RiboZero library prep to deplete cytoplasmic polyadenylated tails. Samples were sequenced to 50 million (M) 2 x 100 bp paired-end reads. Samples were checked for quality using FastQC version 0.11.8 (Andrews, 2010) and aggregated using MultiQC version 0.9 (Ewels et al., 2016). For preprocessing of the RNAseq data, we trimmed adapters using bbduk as part of bbmap version 38.22 (Bushnell, 2014) with the following parameters: `qtrim=rl trimq=30 minlen=75 maq=20`. Post trimming quality was again checked using FastQC version 0.11.8 (Andrews, 2010) and MultiQC version 0.9 (Ewels et al., 2016). Post trimming samples had an average of 35.18M and median of 35M reads (Additional Table 2).

All RNAseq samples were aligned to Gencode GRCh38.p12 human reference genome informed on the sex chromosome complement of the sample (Olney et al., 2020; Webster et al., 2019) using HISAT2 version 2.1.0 for alignment (Kim et al., 2015) and SubRead FeatureCounts version 1.5.2 for quantification (Liao et al., 2014). Briefly, the sex chromosome complement of the sample was first checked by investigating the expression of five Y-linked (*EIF1AY, KDM5D, UTY, DDX3Y, RPS4Y1*) genes and one X-linked gene (*XIST*). A sample with presence of a Y chromosome will show expression for most or all the Y-linked genes while samples with at least two X chromosomes will show expression for *XIST* (Additional Figure 1). Samples with no evidence of a Y chromosome were aligned to a reference genome with the entire Y chromosome masked with Ns to avoid mis-mapping of homologous X-Y sequence reads (Olney et al., 2020). Samples with evidence of a Y chromosome were aligned to a reference genome with the Y chromosome pseudoautosomal regions (PARs) masked out as those regions are replicated 100% on the X chromosome PARs in GRCh38.p12. We followed the XY_RNAseq readme (Olney et al., 2020) to utilize a Ymasked, and YPARs masked reference genome and HISAT `-x index` function to create two sex chromosome complement reference indexes used for alignment. HISAT2 alignment was performed with the following parameters, `--dta` for downstream transcriptome assembly, `--rna-strandness RF` to indicate the sequences are reverse forward, `--phred 33` encoding, and pair-end alignment. RNAseq alignment files were then sorted, read groups were added, duplicates were marked, and files were indexed using bamtools 2.5.1 (Barnett et al., 2011) and Picard 2.9.2 (*Picard Tools - By Broad Institute*, n.d.). FeatureCounts was employed using `--primary` to only use primary alignments and `-p 2` to specify the minimum number of consensus reads from the same pair, suggested for paired-end read data (Liao et al., 2014). FeatureCounts uses the gene annotation file to infer exon-exon junctions from connecting each pair of neighboring exons from the same gene (Liao et al., 2014). FeatureCounts was run twice for each RNAseq sample, once for the gene level using `-g gene_name`. There are 57,133 genes in the Gencode GRCh38.p12 human reference genome used in this analysis.

### Exome data processing

We used FastQC version 0.11.8 (Andrews, 2010) and MultiQC version 0.9 (Ewels et al., 2016) for visualizing quality for whole-exome data. We trimmed adapters using bbduk as part of bbmap version 38.22 (Bushnell, 2014) with the following parameters: `qtrim=rl trimq=30 minlen=75 maq=20`. We used bwa-mem version 0.7.17 (Li, 2013) to align the whole exome samples. Samples were aligned to a sex chromosome complement reference (see RNAseq data processing for more details (Olney et al., 2020; Webster et al., 2019)). Post alignment, PCR duplicates were marked using Picard version 2.18.27 (*Picard Tools - By Broad Institute*, n.d.). To genotype variants, we used GATK version 4.1.0.0 (DePristo et al., 2011; McKenna et al., 2010; Van der Auwera et al., 2013). We first used GATK’s HaplotypeCaller to generate GVCF files. Second, we combined GVCF from 60 individuals, 30 male XY, and 30 female XX, using GATK’s CombineGVCFs.

### Late first trimester placentas

Late first trimester, 10.5 - 13.5 weeks, from last menstrual period when available, human placenta RNAseq samples from Gonzalez et al. 2018 (Gonzalez et al., 2018) were downloaded from NCBI GEO Accession GSE109082 using `fastq-dump -I --split-files` (Leinonen et al., 2011). There are 17 female XX and 22 male XY placenta samples in this data set, all of which self-reported as white (Gonzalez et al., 2018). GSE109082 were processed similarly as the full-term uncomplicated placentas with one exception, trimming for quality. GSE109082 transcriptome samples before trimming had an average of 22.53M 2 x75 bp paired-end reads. GSE109082 paired-end reads were checked for quality using FastQC version 0.11.8 (Andrews, 2010) and MultiQC version 0.9 (Ewels et al., 2016). We preprocessed the samples from GSE109082 similarly to the newly generated samples, with minor changes due to the difference in the RNAseq data: trimmed adapters using bbduk as part of bbmap version 38.22 (Bushnell, 2014) with the parameters `qtrim=rl trimq=25 minlen=40 maq=10`. Post-trimming quality was checked using Fastqc and MultiQC. Post trimming samples had an average of 20.6M and a median of 18.8M reads (Additional Table 2). Reads were then aligned to a sex chromosome complement Gencode GRCh38.p12 reference genome using HISAT2 (Kim et al., 2015). Gene level counts were obtained using SubRead FeatureCounts (Liao et al., 2014). The late first trimester placentas were reverse forward sequencing, the same as the term placentas presented here. Thus, the HISAT2 and FeatureCounts parameters were the same for both the late first trimester (Gonzalez et al., 2018) and the full-term placentas.

### Multidimensional scaling

Multidimensional scaling (MDS) of the expression counts following subRead FeatureCounts was generated using `plotMDS` of the limma package (Law et al., 2014). `plotMDS` is a slightly modified MDS that plots the transcript expression profiles on a two-dimensional scatterplot so that distances on the plot approximate the typical log_2_ fold changes between the samples. MDS plots were generated using the `gene.selection` parameter and selecting “`common`” for all shared genes. This was repeated for the top 100 genes that show the most extensive standard deviations between samples (Additional Figure 2). The full-term placentas were sequenced at two different time points in batch 1 and batch 2. Before clustering with MDS, we accounted for batch effects using the `removeBatchEffect` part of the limma package (Law et al., 2014). This was for visualization purposes only. Batch was included as a covariant in the linear model downstream; see Differential expression.

### Excluding RNAseq samples

Samples that failed quality control (QC) were removed from downstream analysis. Samples were removed that had less than 12.5M or higher than 90M sequences remaining after trimming. If more than 30% of the reads deviated from the sum of the deviations from the normal distribution of the per-sequence GC content as defined by the FASTQC report, then the sample was removed (Additional Table 3). Samples were also excluded that did not cluster with reported sex assigned at birth (Additional Figure 2 & Table 3). There were 23 male XY and 27 female XX full-term placentas that passed QC and were included in the downstream analyses. All 17 female XX and 22 male XY late first trimester placentas from Gonzalez et al. 2018 passed QC and were kept for downstream analysis.

### Subject demographic analysis

We inferred population ancestry from the variants obtained from the whole-exome data using Peddy (Pedersen & Quinlan, 2017) (Additional Figure 3 & Additional Table 1). The resulting outputs of the PCA analysis in Peddy yielded principal components (PC) that were used to assign predicted ancestry. PC1 and PC2 were used later downstream as covariant in the linear model for the differential expression analysis. We did not infer population ancestry from the late first trimester GSE109082 placentas as this data set only included RNAseq data, and not DNA.

### Quantify technical and biological variation in RNAseq expression data

Utilizing variancePartition (Hoffman & Schadt, 2016), a linear mixed model was employed to quantify variation in each expression trait attribute. Variation within gestational age (GA), sequencing lane, sex, reported race, and birth weight was examined. Variation in placenta expression for maternal clinical data, including parity, gravidity, pre-pregnancy body mass index (BMI), and maternal age, were also examined (Additional Figure 4). We did not run variancePartition for all the first trimester placentas because we did not have access to clinical data for this sample set. We additionally examined sex differences for clinical information for full-term placentas for maternal age at delivery, pre-pregnancy BMI, gravidity and parity, gestational age, method of conception, self-reported race, and birth weight. Sex differences for continuous variables were tested using a t-test, p-value < 0.05. A Fisher’s exact test was used to test for sex differences for categorical variables, p-value < 0.05 (Additional Table 4 & Additional Figure 5).

### X and Y gametolog gene expression

A list of X and Y gametologous genes (genes with shared sequence homology and ancestry due to existence as homologs on ancestral autosomes that became sex chromosomes) were curated from a combination of Skaletsky et al. 2003 and Godfrey et al. 2020 (Godfrey et al., 2020; Skaletsky et al., 2003) (Additional Table 5). In samples determined to have a Y chromosome, the CPM value of the X-linked gametolog and the Y-linked gametolog were summed and included in a single value under the X-linked gametolog label. Then we compared expression between the XX female X-linked gametolog expression to XY male X-linked gametolog plus Y-linked gametolog gene expression using a Wilcox rank-sum, p-value < 0.05 (Figure 4, Additional Table 5, Additional Figure 6).

### Differential expression

Sex differential expression analysis between male XY and female XX placentas was performed using the limma/voom pipeline (Law et al., 2014). Quantified read counts from each sample were generated using SubRead featureCounts and combined into a count matrix, with each row representing a unique gene id, and each column representing the gene counts for each unique sample. Using the DGEList function in the limma package the counts matrix and a tab-delimited file containing sample ID, sex, race, batch, lane, GA, parity, maternal age, gravidity, pre-pregnancy BMI, birth weight, PC1 and PC2 from the whole exome data were read into R (Additional Table 1). Technical replicates from within a placenta were summed together using the sumTechReps function in version 3.14.0 (Robinson et al., 2010). Normalization factors were calculated using the calcNormFactors function in EdgeR (Robinson et al., 2010). Lowly expressed genes were filtered out using a minimum threshold of 1 Fragments Per Kilobase Million (FPKM) in at least one group being compared. Then we ran Trimmed Means Method (TMM) to normalize for library size variation between samples (Robinson & Oshlack, 2010). Counts were then transformed to log_2_(CPM+0.25/L), where CPM is counts per million, L is library size, and 0.25 is a prior count to avoid taking the log of zero (Law et al., 2014). For each comparison of interest, a model was created to compare between the groups where each coefficient corresponds to a group mean. The model.matrix function in the ‘stats’ library in R was used to specify the model being fitted. The model was fitted for sex for both the term and first-trimester placenta analyses. Additional covariates (batch, birth weight, lane, and exome sequencing principal components used to account for ancestry) were also added for the term placentas. There were no covariates to add to the model matrix for the late first trimester placentas because clinical and technical information was not made available for this data set. For each differential expression analysis, a linear model was fitted to the DGEList-object, which contained the normalization factors for each gene counts for each sample, using the limma lmfit function which will fit a separate model to the expression values for each gene (Law et al., 2014). Comparisons between groups were then obtained as contrasts of the fitted linear model. An empirical Bayes approach was applied to smooth the standard errors. Genes are defined as being differentially expressed between groups when the adjusted p-value is ≤ 0.05 using a Benjamini-Hochberg false discovery rate (Law et al., 2014) (Figure 1).

**Figure 1.**
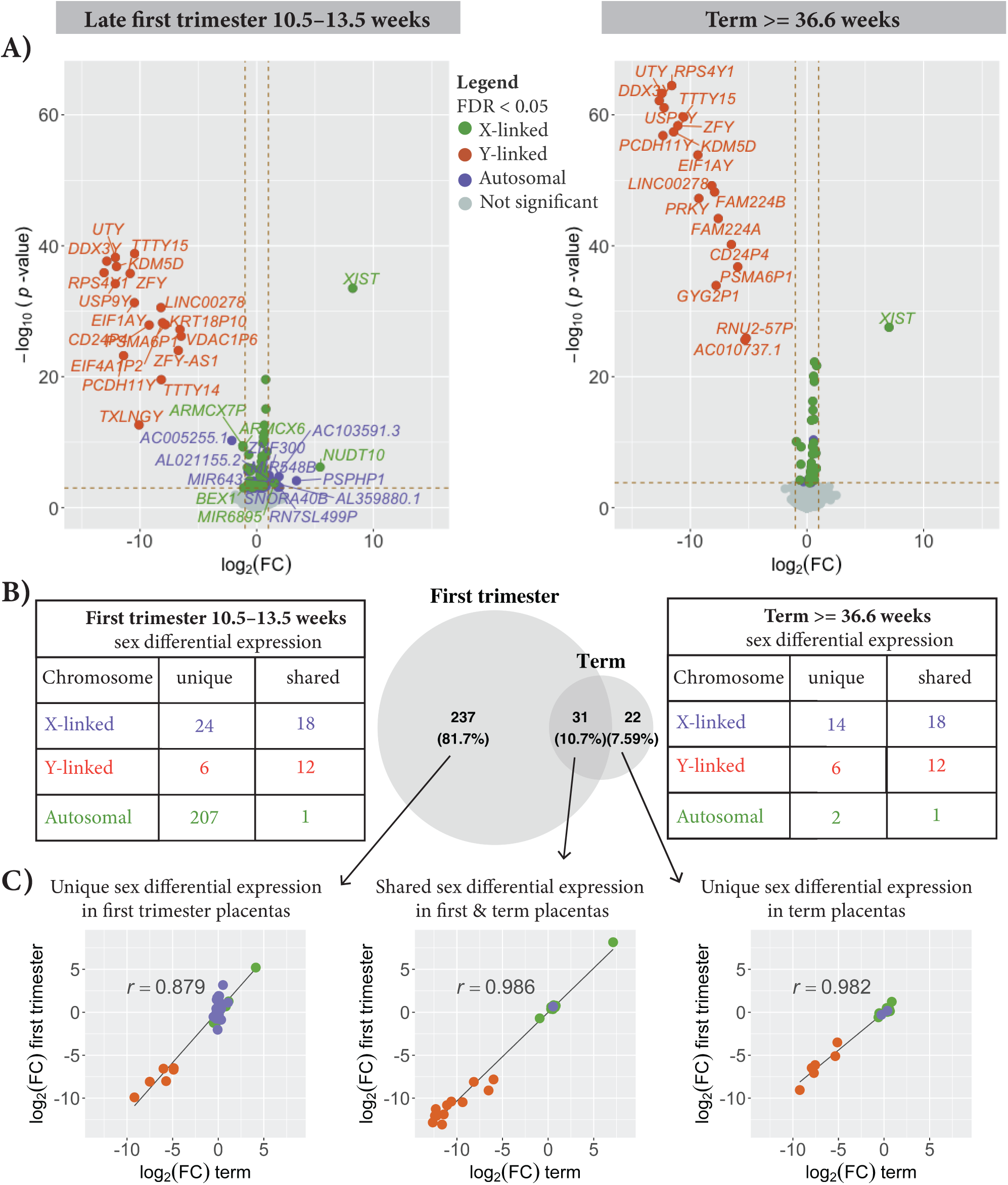
Sex differential gene expression in late first trimester and term placentas are highly correlated. Sex differences in gene expression, log_2_(CPM + 0.25/L), between 17 female and 22 male late first trimester (10.5 - 13.5 weeks) placentas on the left and 27 female and 23 male term ( ≥ 36.6 weeks) placentas shown on the right (A). Each point represents a gene. Genes that are sex differentially expressed, adjusted p-value < 0.05, are indicated in purple for autosomal, orange for Y-linked, and green for X-linked. The number of uniquely sex differentially expressed genes and shared between the late first trimester and the term placentas is shown in (B). More genes are sex differentially expressed in the later first trimester (10.5 - 13.5 weeks) than in the term ( ≥ 36.6 weeks) placentas. There are 237 genes that are uniquely called as sex differentially expressed in the late first trimester placentas that are not called as sex differentially expressed in the term placentas; however, the log_2_ female-to-male expression ratio for those genes are highly correlated between the later first trimester (Y-axis) and the term (X-axis) placentas r = 0.879 (C). There is also a high correlation for the 31 sex differentially expressed genes that are called in both the late first trimester placentas and the term placentas, r = 0.986 and for the 22 genes uniquely called in the term placentas r = 0.982.

### Quantifying sex differences for innate immune gene expression

Differential expression between male XY and female XX placentas for 979 innate immune genes, as defined by InnateDB (Breuer et al., 2012) (Additional Table 6). The InnateDB is a publicly available database of genes, proteins, experimentally verified interactions, and signaling pathways involved in the innate immune response to microbial infection. Sex differential expression of the 979 innate immune genes was performed using the limma/voom pipeline (Law et al., 2014). We repeated this analysis for both the term and late first trimester placentas. The model matrix for term placentas was the same when looking at the differential expression for the whole transcriptome. Only genes determined to be expressed in at least one sex were included in the analysis (see Methods). In addition to differential expression, we generated an MDS plot on the expression data for only the innate immune genes to determine immune gene expression similarity among samples (Figure 2).

**Figure 2.**
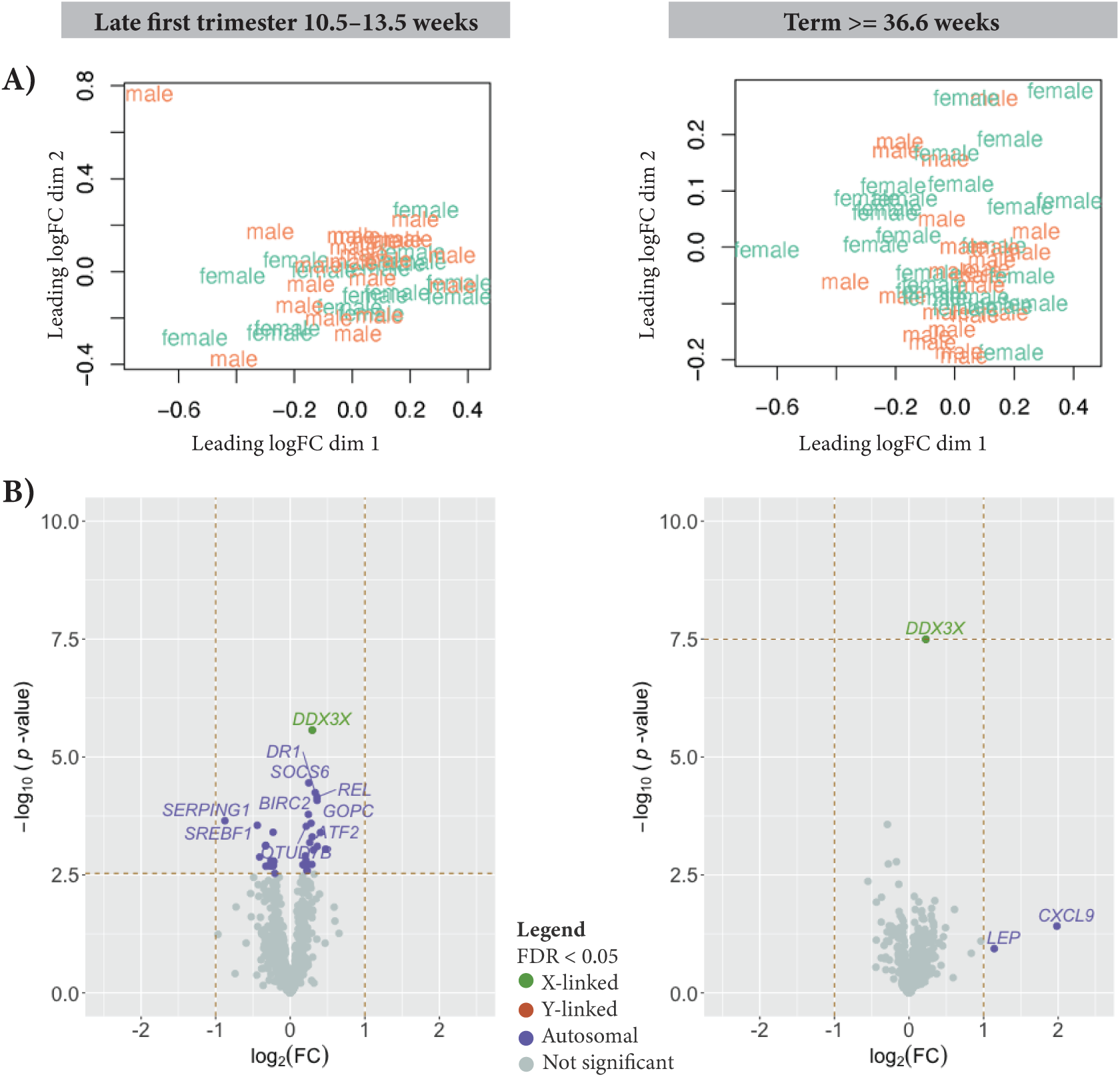
Innate immune gene expression between male XY and female XX placentas. Of the 979 innate immune genes from InnateDB, 625 genes are expressed in the late first trimester placentas and 628 are expressed in the term placentas. (A) MDS plot shows no clustering by genetic sex in either the late first trimester (left) or term (right) placentas. (B) volcano plot of the sex differential expression for late first trimester placentas (left) and term placentas (right). Each point represents a gene. Genes that are sex differentially expressed, adjusted p-value < 0.05, are indicated in purple for autosomal, orange for Y-linked, and green for X-linked.

### Gene function and enrichment network analysis

We used the TopFunn webtool to infer the function of genes and identify enriched biological processes, molecular functions, and gene families (J. Chen et al., 2009). Additionally, we looked at each sex differentially expressed gene from the late first trimester and full-term placenta comparisons using genecards.org and a literature review of genome-wide association studies and expression quantitative trait loci (eQTL) to investigate if that gene is involved in known diseases or disorders, particularly with known pregnancy complications (Additional Table 12).

### Sex differences in adult GTEx tissues

For each of the 42 non-reproductive adult tissues in the Genotype by Tissue Expression (GTEx) consortium, we computed the log_2_ female-to-male expression ratios from the reported Transcripts Per Kilobase Million (TPM) counts version 2017-06-06_v8 (Additional Table 8) (Carithers et al., 2015; Oliva et al., 2020). To determine if sex differences in gene expression within the placenta are correlated with sex differences in adult tissues, we computed the coefficient of correlation, r, of the log_2_ female-to-male expression ratios for sex differentially expressed genes found in the placenta to the log_2_ female-to-male expression ratios of the same genes in each adult tissue (Figure 3).

**Figure 3.**
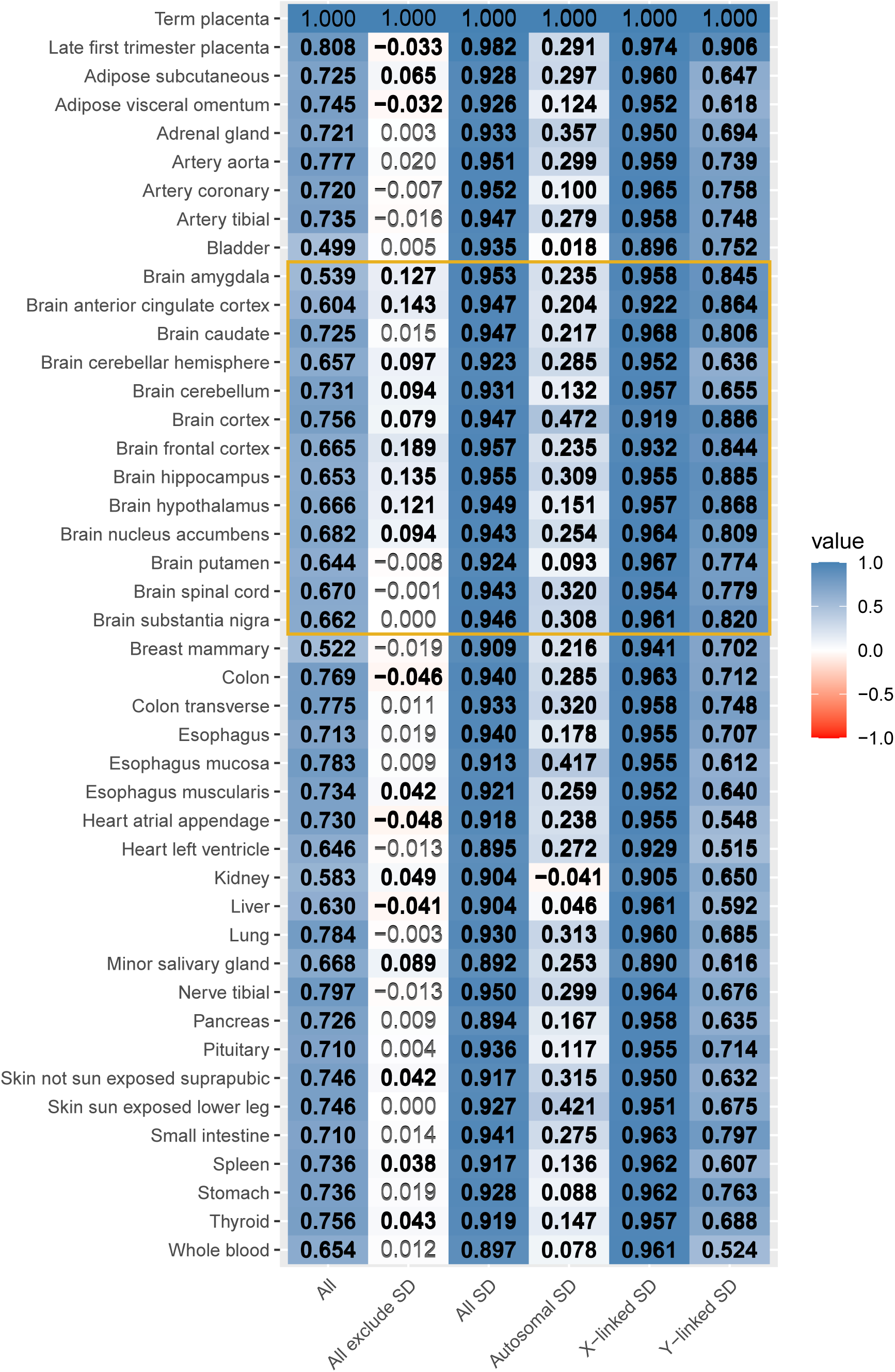
Correlation of female-to-male expression between placenta and 42 non-reproductive adult tissues. From left to right: The first column is All genes which includes 11,179 expressed genes in the placenta and contains count information in GTEx. Of the 11,179 genes 10,762 are autosomal or MT genes, and 417 are sex-linked (X or Y-linked). The second column is All genes but excluding sex differentially (SD) genes. Exclude SD genes comprises 10,936 genes, of which, 10,580 are autosomal or MT genes, and the remaining 356 are X-linked. All SD genes includes 243 sex differentially expressed genes in the placenta (late first trimester or term) adjusted p-value < 0.05 and contains count information for GTEx tissues. Of the 243 sex differentially expressed genes, 182 are autosomal and 61 are sex-linked (49 X-linked and 12 Y-linked). The last three columns show the coefficient correlation, r, in the log_2_ female-to-male expression ratios between term placenta, late first trimester placentas, and 42 non-reproductive adult GTEx tissues when only exaiming the autosomal sex differentially expressed genes (182 genes), SD and X-linked (49 genes) and SD and Y-linked (12 genes). The gold box highlights adult samples from many regions of the brain which all showed high correlation of with differential expressed sex-linked genes in the placenta. Black and bold indicates a significant correlation after correcting for multiple testing, adjusted p-value < 0.05.

## Results

### Unsupervised separation of placenta samples by sex

Multidimensional Scaling (MDS) analysis was performed on full-term and late first trimester placentas to determine trends in the placenta transcriptome. MDS of the full-term placentas show separation of placenta by genetic sex on the first dimension showing that sex is a key factor in gene expression in the placenta. Each placenta sample from term placentas included two replicate samples, all passed our QC except one female XX placenta replicate, so both replicates for this sample was removed from the downstream analyses. The replicate failed QC due to a low number of sequence reads after filtering and falling outside of the expected MDS cluster (Additional Figure 2 and Additional Table 3).

### Population ancestry inferred from whole exome data

Principal component analysis of the full-term placenta whole exome data shows the samples separated by reported and inferred ancestry in many cases (Additional Table 1). The self-reported race and ethnicity for the placenta samples included 14 Asian, 20 Black, 4 Hispanic, 20 White, and 2 unknown (Additional Table 1). The ancestry prediction estimates that the population ancestry of the samples is: 14 Asian (5 South Asian, 3 East Asian, 1 European, and 2 unknown), 20 Black (17 African, 3 unknown), 4 Hispanic (1 European, 3 unknown), 20 white (15 European, 1 American, 1 South Asian, 1 African, 2 unknown) and 2 unknown (1 American, 1 South Asian). To account for population ancestry differences among the samples, PC1 and PC2 from the whole exome data was included as covariates in the linear model used to identify sex differential expression.

### Modeling parameters to determine sex differences in placental gene expression

To determine differentially expressed genes in the placenta, specifically because of differences in infant sex, we assessed non-sex variables that show differences between the sample groups that could confound our results. First, we examined differences in clinical phenotypes collected in this study. Nearly all the full-term placentas collected for this study were spontaneous methods of conception (all but one male from an IVF pregnancy and one female placenta from intrauterine insemination) (Additional Table 1) making this an unlikely confounding variable for sex differences in gene expression. Maternal age at delivery ranged from age 22 to 45 years old, and pre-pregnancy BMI ranged from 19.40 to 66.30. There was no significant difference in maternal age or pre-pregnancy body mass index (BMI) for women who carried a male XY versus women who carried a female XX (t-test p-values = 0.39, and 0.73, respectively). Gravidity, the number of pregnancies the mother had achieved, ranged from 1 to 9, and parity, the number of pregnancies reaching higher than 20 weeks, ranged from 0 to 4; the current pregnancy at the moment the data was collected was not included in the parity counts. Gravidity and parity did not show a significant difference in women that carried a male XY or female XX pregnancy (t-test p-values of 0.43 and 0.61, respectively). Gestational age ranged from 36.6 to 41.1 weeks with no significant difference between people who carried a male XY versus a female XX pregnancy (t-test p-value = 0.90). The difference in average birth weight approached statistical significance between the sexes, with a male mean of 3.6 kilograms in males and female mean of 3.3 kilograms (t-test p-value = 0.056; Additional Figure 5 & Additional Table 4). Since a previous study demonstrated sex differences in birth weight and placental genes differentially expressed by birth weight (Crawford et al., 1987; Sõber et al., 2015), we included birth weight as a covariate in our model. Factoring out the birth weight by including it as a covariate allowed us to focus our results on genes differentially expressed solely based on genetic sex, as this is what our study was designed to assess. If birth weight was not included in the model as a covariate, we identify 19 more genes as differentially expressed, only one of which, ASMTL, was also differentially expressed in the late first trimester placentas (Additional Table7). In addition to assessing differences in clinical variables between sexes, we used variance partitioning to assess the extent to which clinical features as well as technical features explained the variance in gene expression between our placenta samples. Variance partitioning was performed for the full-term placenta RNAseq samples revealing that the sequencing lane explained the largest amount of variance in our data (see methods; Additional Figure 4). Maternal variables tested explained very little of the variance in placenta gene expression, consistent with the same finding reported in a previous placenta gene expression study (Sõber et al, 2015). Our term placenta samples were collected in two batches, so the batch was also included as a covariate. Using results from all of these approaches, we constructed a linear model to determine sex differentially expressed genes that included batch, lane, the first principal components from exome sequencing which captured differences in population ancestry, and birth weight as covariates.

### Sex differential expression from male XY and female XX term uncomplicated human placentas

We observed 14,441 genes expressed with an FPKM > 1 in at least all the male XY or all female XX placenta samples (Additional Table 9). 53 genes exhibited sex differentially expressed with an adjusted p-value < 0.05 (Figure 1 & Additional Table 7). Thirty genes showed higher expression in the female XX placentas and 23 genes showed higher expression in the male XY placentas. Of the 30 genes that were more highly expressed in the female XX placentas than the male placentas, 28 are X-linked and two are autosomal (EIF2S3B, and EIF1AXP1). Of the 23 genes more highly expressed in males, 18 are Y-linked, 4 are X-linked (CD99, RPS6KA6, VDAC1P1, VAMP7), and one is autosomal (PRKCE) (Figure 1 & Additional Table 7).

### Sex differential expression within late first trimester placentas

There were 13,502 genes expressed with an FPKM > 1 in at least the male XY or female XX late first-trimester placenta samples that were reprocessed here using the same approach as the term placentas (see Methods) (Gonzalez et al., 2018). After reprocessing the sequencing data to improve alignment to sex chromosomes, we identified 268 genes with a differential expression between male and female placentas with an adjusted p-value < 0.05 (Figure 1). Of these, 180 genes showed higher expression in the female XX placentas and 88 genes showed higher expression in the male XY placentas.

### Sex differential expression shared between late first trimester and full-term placentas

We observed that sex differences in gene expression ratios between female and male placentas were highly correlated between late first trimester and term placentas. The late first trimester (Gonzalez et al., 2018) and full-term placenta RNAseq samples were processed using the same tools and only differed in the trimming parameters (bbduk minlen, see Methods) and the covariates added to the linear model for computing sex differential expression (see above). Of the 268 genes that were sex differentially expressed in the late first trimester placentas, 31 or 10.7% were shared with the genes identified as sex differentially expressed in the full-term placentas (Figure 1). Although there were more genes identified as sex differentially expressed in the late first trimester placentas given our adjusted p-value threshold < 0.05, the log_2_ female-to-male expression ratio for these genes was highly correlated between late first trimester and term placentas (Figure 1C). For the 237 genes that were uniquely called sex differentially expressed in the late first trimester placentas, the correlation coefficient, r, for the log_2_ female-to-male expression ratio between late first trimester and term placentas was 0.879. The correlation coefficient r for the log_2_ female-to-male expression ratio between the late first trimester and term placentas for the 31 genes differentially expressed in both placenta datasets was 0.986. Twenty-two genes were uniquely identified as sex differentially expressed in the term placentas compared to the late first trimester placentas; with the correlation coefficient of log_2_ female-to-male expression ratio for these genes being 0.982.

### Biological processes and molecular functions enriched in placenta of a specific sex

We investigated enrichment of biological functions and processes in genes that were overexpressed in one sex versus the other sex for placentas collected from full-term uncomplicated pregnancies and late first trimester (Gonzalez et al., 2018) (adjusted p-value < 0.05) (Additional Table 7, Additional Table 11). Genes more highly expressed in male XY term placentas than female XX term placentas were found to be involved in histone lysine and protein demethylation processes and histone demethylase activity, driven by Y-linked genes, including UTY and KDM5D (previously also called SMCY or JARID1D). Serine/threonine kinases RPS6KA6 (chrX), PRKCE (chr2), and PRKY (chrY) were also more highly expressed in term male XY placentas. Five genes expressed exclusively in term male XY placentas were identified as members of minor histocompatibility antigen gene family: UTY, DDX3Y, RPS4Y1, KDM5D, and USP9Y (all Y-linked genes). In contrast, genes more highly expressed in female XX term placentas than in male XY term placentas were identified as involved in translational initiation and regulation of sister chromatid cohesion, driven mainly by X-linked genes including RPS4X, DDX3X, NAA10, EIF1AX, EIF2S3, and HDAC8. In the late first trimester placentas, as in term placentas, genes more highly expressed in male XY placentas compared to female placentas also included minor histocompatibility antigen genes. Genes more highly expressed in the female late first trimester placentas than male late first trimester placentas were shown to be involved in diphosphatase and phosphotransferase activity driven by DCP2, NUDT10, NUDT14, PIGF, and PIGN and branched-chain amino acid catabolic and metabolic processes driven by ACADSB, DBT, HSD17B10, ALDH6A1. In both term and late first trimester placentas, about 30% of the sex differentially expressed genes were associated with pregnancy complications (Additional Table 12). Miscarriage and preeclampsia were the most frequently observed complications, but there were others, such as gestational diabetes and fetal growth restriction.

### More sex differences in expression of immune and immune modulator genes in late first trimester placentas than in term placentas

We tested for sex differences in a subset of genes with known connections to innate immune function in this RNAseq study of late first trimester and term placentas. Of the 979 innate immune genes reported from InnateDB, 628 were expressed in the term placentas. Unlike with all genes where placenta samples clearly clustered by sex (Additional Figure 2), multidimensional scaling with only the innate immune genes showed no distinguishable pattern (Figure 2). Of the 628 innate immune genes expressed in the term placentas, only DDX3X showed a statistically significant difference in expression (adjusted p-value = 2E-5; Figure 2; Additional Table 6). This result was further investigated in the gametolog section as DDX3X has a functional Y-linked copy, DDX3Y.

Unlike in the term placentas where only one innate immune gene showed differential expression by sex, the late first trimester placentas showed 37 innate immune genes as differentially expressed between the sexes (adjusted p-value < 0.05; Additional Table 6). Of the 979 innate immune genes reported from InnateDB, 626 were expressed in the late first trimester placentas (see Methods). Like the term placentas, DDX3X also showed a difference in expression (See Gametolog section; Additional Tables 5 & 6). SERPING1 showed the highest fold change, with 1.84 times greater average expression in late first trimester male XY placentas than in female XX placentas (Figure 2). Seven of the differentially expressed innate immune genes had transcription activator activity (SREBF1, REL, RELA, CREB1, ATF2, DDIT3, and STAT6), seven had kinase activity (PIK3CA, SOCS6, SOCS5, TRIM28, GSK3A, DDX3X, RBCK1), and seven had ubiquitin transferase activity (BIRC2, TRIM13, TRIM28, HERC5, RBCK1, HACE1).

### Female-to-male gene expression ratios in the placenta are correlated with adult tissues

Sex differences in gene expression in the human placenta were highly positively correlated with sex differences in adult tissues, and this was largely driven by sex-linked genes (Figure 1, 3, & Additional Table 7). We first observed that all genes expressed in the placenta showed a significant correlation between the log_2_ ratio of female-to-male expression in the placenta and adult tissues (“All” in Figure 3). However, when we removed sex-differentially expressed genes, this correlation was markedly decreased (“Exclude SD” in Figure 3). Conversely, we observed high correlation between the subset of sex-differentially expressed genes in the placenta and 42 adult tissues, with a pairwise r ranging from 0.892 to 0.982 (adjusted p-value < 0.05 for all comparisons; “All SD” in Figure 3). The adult tissue with the highest correlation to that of term placentas in the log_2_ female-to-male expression was the frontal brain cortex with an r of 0.957 (and adjusted p-value < 0.05). Notably, the log_2_(female-to-male expression ratio) correlation for placenta sex differentially expressed genes between term placentas and adult tissue brain regions ranged from r of 0.923 to 0.957(adjusted p-value < 0.05; “All SD” in Figure 3).

The high r of the log_2_female-to-male expression between term placenta to adult tissues for genes found to be sex differentially expressed in the placenta was largely driven by sex-linked genes (Figure 3). Of the 243 sex differentially expressed genes in the placenta and adult GTEx tissues, 182 were not sex-linked (autosomal or mtDNA), and the remaining 61 were on the sex chromosomes, X or Y (Additional Table 7). Despite showing sex differences in the placenta, the autosomal genes showed much lower, and largely not significant, correlations between placenta and adult tissues compared to the sex-differentially expressed genes (“Autosomal SD” in Figure 3). In contrast, we generally observed equal or higher correlations (r ranges from 0.890 to 0.967, adjusted p-value < 0.05) between the placenta and adult tissues for X-linked genes compared to any other group of sex-differentially expressed genes—again with the highest correlations between the placenta and adult brain tissues (“X-linked SD” in Figure 3). Y-linked genes showed an intermediate range of correlation values (r from 0.515 to 0.886; “Y-linked SD” in Figure 3), but all were significant after correction for multiple testing, unlike autosomal sex-differentially expressed genes.

### Sex differences in expression for X-linked gametolog genes

X-linked gametolog genes showed a difference in expression between female and male placentas, but the bias of expression differed whether the Y-linked gametolog was included in the analysis (Figure 4). Of the 23 gametologous gene pairs (genes with a functional copy on the X chromosome and Y chromosome due to shared ancestry on the sex chromosomes), 14 were expressed in the late first trimester and term placenta samples (FPKM > 1 in at least all the male or all the female samples; Additional Table 9). In the late first trimester placentas, 7 out of the 14 X-linked gametolog genes showed higher expression in females compared to males (DDX3X, ZFX, KDM6A, PRKX, PRS4K, EIF1AX; Additional Table 5). When we summed the expression of the X and Y-linked gametolog for male samples, we no longer saw a sex difference in expression for ZFX, PRKX, and EIF1AX, Wilcoxon test p-value > 0.05 (Additional Table 5). Three sex differentially expressed genes, DDX3X, KDM5C, and RPS4X, continued to show higher expression in female XX compared to male XY late first trimester placentas. On the other hand, when we took the sum X + Y-linked expression in males, KDM6A flipped direction and showed higher expression in male compared to female late first-trimester placentas, Wilcoxon test p-value < 0.01 (Additional Table 5). Additionally, PCDH11X changed from showing no sex difference in expression to showing higher male expression compared to female in late first trimester placentas, Wilcoxon test p-value < 0.01 (Figure 4 & Additional Table 5).

**Figure 4.**
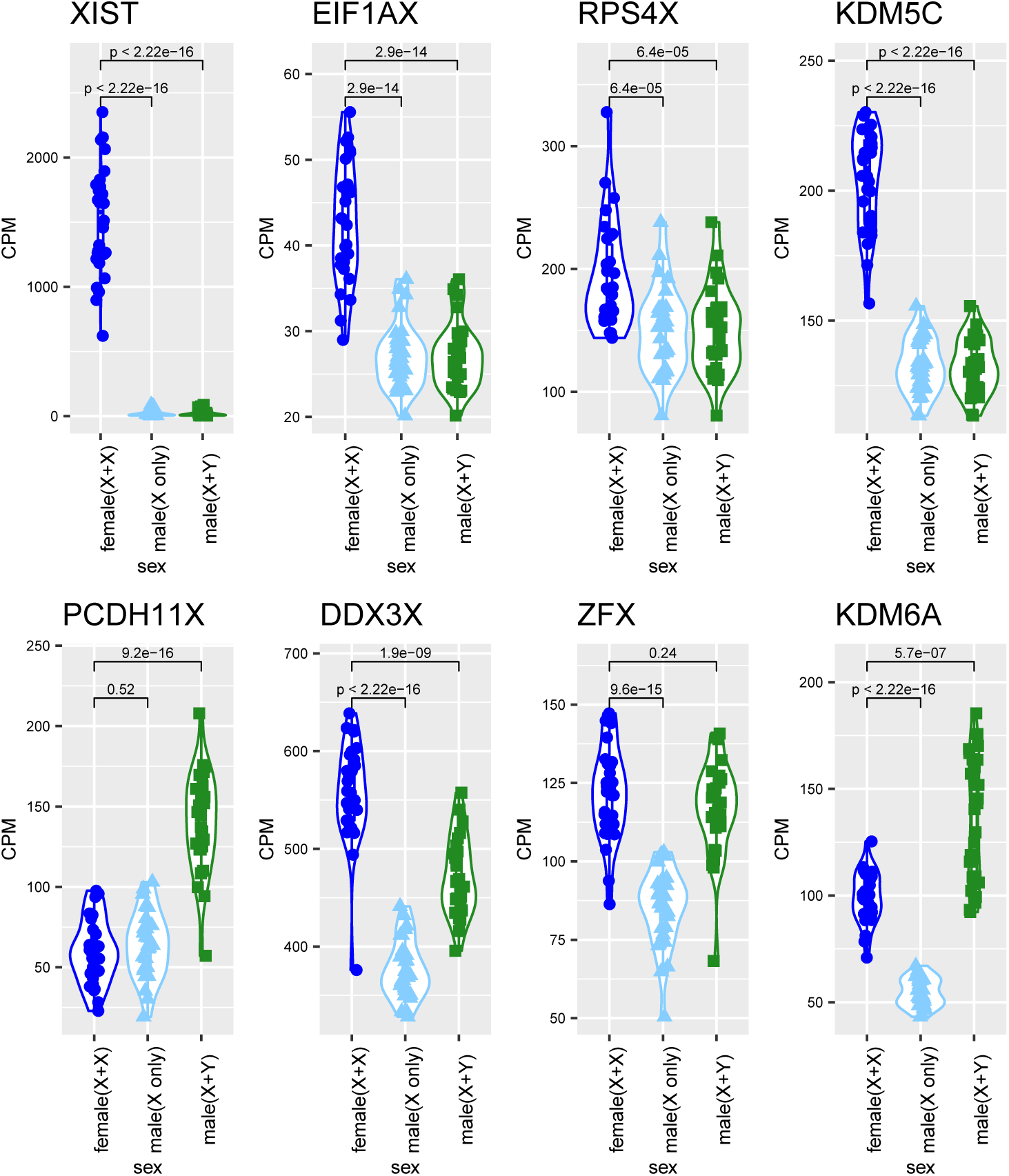
Chromosome X+Y expression of gametologs in XY male placentas. There is a significant difference in male XY to female XX expression for ZFX and KDM6A (UTX) when only looking at the X chromosome CPM expression value. When we add the Y chromosome-linked CPM expression count for these genes for male samples, there is no longer a difference in expression between males XY and females XX for ZFX. KDM6A changes directionality and shows males as having significantly higher expression than females when accounting for gametolog expression. PCDH11X, when adding Y-linked CPM expression, shows a significantly higher expression than females. T-test to see if there is a difference between the female CPM and the male CPM for each gene, p-value < 0.05.

In term placentas, we observed a similar pattern when we summed the X and Y-linked expression for males. The same 7 X-linked gametolog genes that showed higher expression in the female late first-trimester placentas also showed higher expression in the female term placentas compared to male placentas (Additional Table 5, Additional Figure 6). When we took the sum expression of the X and Y-linked copy for male XY term placentas, we no longer saw a sex difference in expression for ZFX, PRKX, and RPS4X, p-value > 0.05 (Additional Table 5). This differed from the late first trimester placentas that showed sex higher expression in female placentas for RPS4X, regardless of accounting for the Y-linked gametolog (Additional Table 5). Three genes, DDX3X, KDM5C, and EIF1AX, continued to show higher expression in female compared to male XY term placentas compared to female XX placentas after accounting for Y-linked gametolog expression, Wilcoxon p-value < 0.01 (Additional Table 5). Similar to late first trimester placentas, when we accounted for Y-gametolog expression, KDM6A flipped direction and showed higher expression in male compared to female term placentas and PCDH11X changed from showing no sex difference in expression to showing higher expression in male compared to female term placentas, Wilcoxon test p-value < 0.01 (Figure 4 & Additional Table 5).

## Discussion

### Sex differences in gene expression begin in early pregnancy and continue through adulthood

We observed a positive correlation in the log_2_ female-to-male expression ratio for sex differentially expressed genes between the late first trimester (Gonzalez et al., 2018) and term placentas, and between term placentas and 42 non-reproductive adult GTEx tissues (Figure 3). A previous study using microarrays compared first trimester and term placentas and found thousands of genes to be differentially expressed, but they did not characterize sex differences (Sitras et al., 2012). A previous microarray study on placental cell types reported 11 genes that were sex differentially expressed in all trophoblast and endothelial cell types, all of which are sex differentially expressed in the late first trimester and term placenta samples analyzed here (Cvitic et al., 2013). Another study of term placentas reported only X- and Y-linked genes as differentially expressed between female and male placentas (Sõber et al, 2015), consistent with our results which revealed the majority of sex differentially expressed genes on chromosomes X and Y. When we looked at all genes located on the sex chromosomes, X & Y, the log_2_ female-to-male expression ratio was positively correlated between late first trimester and term placentas, as well as adult tissues (Figure 3). When this was repeated for only autosomal genes, 1-22 and mitochondrial chromosomes, we did not observe the same positive correlation between late first trimester, term placentas, and adult tissues (Figure 3). These findings suggest that sex differences in gene expression for sex-linked genes develop early in embryonic tissue and are replicated in adult tissues. Sex differential expression for autosomal genes may be more tissue-dependent, as previously suggested by Lopes-Ramos et al. 2020. Lopes-Ramos et al. 2020 found that sex differentially expressed genes common among adult tissues were enriched for sex chromosome genes, and sex differences for autosomal genes were tissue-specific (Lopes-Ramos et al., 2020). In summary, we observed a positive and significant correlation in the log_2_ female-to-male expression ratio for sex-linked genes between term placentas and adult tissues (Figure 3).

Our study showed that sex differences in placental gene expression is correlated to sex differences in adult tissues, particularly in the brain. In addition to hormones, the placenta produces a host of neurotransmitters to facilitate fetal brain formation (Rosenfeld, 2021). Dopamine and norepinephrine are expressed in placenta cytotrophoblasts in early pregnancy and syncytiotrophoblast cells in late pregnancy (Zhu et al., 2002). This normal expression was altered in pregnancy complications such as pregnancy-induced hypertension syndrome and developmental exposure to endocrine-disrupting chemicals such as bisphenol A and S (Mao et al., 2020; Zhu et al., 2002). Sex differences in fetal brain development have been associated with sex-linked genes such as O-linked N-acetylglucosamine transferase (OGT), which is expressed at a higher level in XX females because it escapes X chromosome inactivation, leading to increased histone methylation and decreased gene expression thought to provide a neuroprotective state in females (Bale, 2016). Taken together, sex differentially expressed genes in the placenta that are correlated to sex differentially expressed genes in the adult brain could provide insight into mechanisms driving brain development differently in each sex and potentially how these mechanisms are altered in neurological disorders.

### Gene enrichment of sexually dimorphic genes reveals genes that may be involved in pregnancy complications

Sex differentially expressed genes may be involved in biological pathways related to pregnancy complications. In term placentas, genes upregulated in XY males are involved in histone lysine and protein demethylation processes and histone demethylase activity, driven by Y-linked genes, including UTY and KDM5D (Additional Table 11). A review of ruminant placenta gene targeting found histone lysine demethylase 1A and androgen signaling involved in gene networks for cell proliferation and angiogenesis (Hord et al., 2020). The authors also note previous studies that have examined exposure to testosterone during pregnancy leading to ovarian dysfunction and low-birth-weight for female offspring suggesting that increased androgen signaling dysregulates fetal development for female offspring (Hord et al., 2020). Five genes in the minor histocompatibility antigen gene family were also upregulated in term male placentas. It has been shown that minor histocompatibility antigens are involved with maternal immune exposure to fetal antigens needed to prevent fetus rejection (Linscheid & Petroff, 2013). Genes upregulated in XX female term placentas are involved in translational initiation and regulation of sister chromatid cohesion, driven mainly by X-linked genes, including NAA10. NAA10 is involved in post-translational protein modifications, and mutations in NAA10 are known to cause Ogden syndrome, which may lead to growth failure (Lee et al., 2017) (Additional Table 11). In a Naa10 mouse knockout study, the authors reported placental insufficiency that contributed to embryonic and neonatal lethality (Lee et al., 2017). Naa10 mouse knockouts showed low birth weight and postnatal growth failure compared to control mice (Lee et al., 2017). Loss of NAA10 plays a role in the development of cardiovascular and growth defects in humans and mice (Lee et al., 2017; Wu & Lyon, 2018). In our study, NAA10 is upregulated in females compared to male term uncomplicated placentas (Additional Table 7).

Using novel sex-informed alignment and functional enrichment analysis, we reanalyzed the Gonzalez et al. 2018 late first-trimester placenta data to compare to our newly collected full-term placenta data. We found genes upregulated in male XY compared to female XX late first trimester placentas are involved in positive regulation of anoikis (a form of programmed cell death). We found that genes upregulated in the female late first trimester placentas are involved in organonitrogen compound catabolic, branched-chain amino acid catabolic, and positive regulation of protein K63-linked ubiquitination processes. NUDT10 was shown to be enriched in these biological processes and is upregulated in female XX late first trimester placentas and was reported in the Gonzalez et al. 2018 study as well (Gonzalez et al., 2018). The role of NUD10 in placenta function remains to be further explored. Additionally, of the 58 genes previously identified as sex differentially expressed in the late first-trimester placentas by Gonzalez et al. 2018, we found 45 of those genes to be sex differentially expressed in the samples using different tools to process the data. Overall, we replicated the findings from Gonzalez et al. 2018 and identified 210 additional genes to be sex differentially expressed among the late first trimester placentas, several of which have been associated with pregnancy complications (Additional Table 10 and 12). We further observed a high positive correlation in the log_2_ female-to-male expression ratio for all 268 sex differentially expressed in the late first trimester placentas compared to that of the term placentas (Figure 1).

### Sex differences in immune gene expression linked to pregnancy complications

The placenta is an immune modulator in the uterine environment interacting with the maternal decidua cells to promote an immunosuppressive environment for maintaining fetal tolerance (Mor et al., 2011; PrabhuDas et al., 2015; Xin et al., 2014). The placenta promotes inflammation response with up-regulation of pro-inflammatory cytokines during early implantation and late in pregnancy (Mor et al., 2011; PrabhuDas et al., 2015). Placentas from preeclampsia pregnancies have been reported to show lower expression of immune protein CD74 and enrichment for the IL-1-signaling pathway compared to uncomplicated placentas (Przybyl et al., 2016).

Furthermore, differences in maternal immune function are different based on the sex of the fetus, such as maternal cytokine production increases throughout pregnancy and are higher when carrying a female fetus (Mitchell et al., 2017). Immune gene expression within the placenta plays a role in maintaining a pregnancy to term (PrabhuDas et al., 2015; Xin et al., 2014); we, therefore, sought to characterize sex differences in immune gene expression in the late first trimester (Gonzalez et al., 2018) and term uncomplicated placentas to expand on previously reported sex differences among uncomplicated placentas (Gonzalez et al. 2018 and Sood et al. 2006).

While an inflammatory response has been shown to happen in the first and third trimesters of pregnancy (Mor et al., 2011), we observed sex differences in immune genes almost exclusively in late first trimester placentas. In the term placentas, when looking at sex differences for only innate immune genes, only DDX3X showed a difference in expression between the sexes, with higher expression in female placentas compared to males (adjusted p-value = 5.55 x 10^-8^) (Figure 2). DDX3X is essential in cell cycle control, and loss of Ddx3x in male mice resulted in early post-implantation lethality (C.-Y. Chen et al., 2016). In female mice, inactivation of a paternal Ddx3x copy resulted in placental abnormalities and embryonic lethality (C.-Y. Chen et al., 2016). This suggests that the expression of DDX3X in the placenta may be critical for proper placental development. DDX3X may show higher expression in female than male placentas because DDX3X escapes X chromosome inactivation in female uncomplicated placentas, showing expression of both the maternal and paternal gene copy (Phung et al., n.d.). Although not significantly differentially expressed between the sexes, there were several immune genes that showed higher mean expression in female or male placentas. CXCR4 showed 1.5 fold higher expression in term male placentas (p-value = 0.004). CXCR4 is a chemokine receptor shown to be expressed in both early and term placenta (Kumar et al., 2004) and shown to have higher expression in mothers with preeclampsia and gestational hypertension (Zheng et al., 2021). CXCR4 showed sex-dependent overexpression in lung adenocarcinomas and the signal was shown to be higher in premenopausal women compared to postmenopausal women and men (Rodriguez-Lara et al., 2014). AQP3 showed a 2 fold higher mean expression in term female placentas. AQP3 is an aquaporin water channel cell membrane protein expressed in both placenta and fetal membranes that has been suggested to be involved in regulation of amniotic fluid homeostasis (Szpilbarg & Damiano, 2017). Cell line studies showed that the inhibition of AQP3 reduces cell migration in trophoblast cells (Alejandra et al., 2018) and expression of AQP3 was found to be significantly reduced in preeclampsia (Szpilbarg & Damiano, 2017). Sex differences in expression of other members of the aquaporin family have been observed in other organs in rodent models (Fan et al., 2005; Nicchia et al., 2001; Sun et al., 2007), but sex differences in placenta expression has yet to be studied. Overall, except for DDX3X, we observed a lack of sex differences in gene expression of innate immune genes among uncomplicated term placentas (Figure 2 & Additional Table 6), suggesting expression of these genes may be important for maintaining all pregnancies to term regardless of the sex of the fetus.

In the late first trimester placentas (Gonzalez et al., 2018), 37 innate immune genes were differentially expressed between the sexes (adjusted p-value < 0.05), including DDX3X (Additional Table 6). Of the 37 innate immune genes differentially expressed between the sexes, SERPING1 showed the highest expression ratio between male to female placentas (log_2_(fold change=1.7). SERPING1 encodes for a highly glycosylated protein and is involved in inhibiting C1r and C1a of the complement component. SERPING1 may be involved in the placental circulatory function, the dyssregulation of which could lead to vascular leakiness and subsequent edema—which often accompany preeclampsia (Vaiman et al., 2005). Intriguingly, unmethylated CpG islands by the transcription start site of SERPING1 have been associated with eight times higher gene expression in placenta tissue from patients with preeclampsia (Blanch et al., 2003; Chelbi et al., 2007). Similarly, SREBF1, which also showed significantly higher expression in males, showed lower methylation rates and higher expression in placentas in pregnancies after using assisted reproductive technologies (Lou et al., 2014), suggesting potential links to uterine conditions that hinder implantation. Overall, we observed many genes that showed sex differences in expression for innate immune genes in the late first trimester placentas, but not in the term placentas, though some of this may have been due to our ability to account for covariates in the analysis of term placentas.

### Quantification of X-linked gametolog genes

Measuring gene expression on sex chromosomes is complicated due to areas of highly conserved regions between the X and Y chromosomes (Olney et al., 2020). Quantifying gene expression is particularly problematic for gametolog genes which have functional copies on both the X and Y chromosomes that are thought to be expressed off of both X and Y chromosomes in males (Wilson & Makova, 2009). For genes present only on the X chromosome, silencing of one X in XX females is thought to equalize expression of the one X in XY males. However, some genes escape X inactivation in XX females and some gametologs show expression from the Y chromosomes in XY males making functional quantification of these genes more difficult (Godfrey et al., 2020; Wilson Sayres & Makova, 2013). In this study we assessed how summing expression of gametologs with X- and Y-linked expression in XY males may alter how we interpret sex differences in gene expression, in contrast with comparing only X-linked gametolog expression. Gene expression differences between female and male placentas in both the full-term placentas (Figure 4, Additional Table 5) and late first trimester placentas (Additional Figure 6, Additional Table 5) for DDX3X, ZFX, PCDH11AX, and KDM6A (UTX) would all be interpreted very differently if expression from X- and Y-chromosomes would have been summed in XY males. For all 4 of these genes, previous literature has demonstrated that they are transcribed from both the X- and Y-chromosomes and that the gametologous genes have similar functions. DDX3X and DDX3Y share over 90% sequence similarity and were observed to have redundant roles in protein synthesis (Venkataramanan et al., 2021). KDM6A (also called UTX) has been shown to escape X-inactivation, and mouse model studies of Y-linked gametolog UTY showed that it maintains some of its function (Gažová et al., 2019). For the same reasons, ZFX and ZFY and protocadherin PCDH11X and PCDH11Y expression were both previously summed in cell line studies that demonstrate dosage compensation on genes from the sex chromosomes (Johansson et al., 2016; Schneider-Gädicke et al., 1989). Properly quantifying gene expression of gametologs from the sex chromosomes can help us to more accurately identify sex differences in gene expression.

## Perspectives and Significance

Overall, these results highlight the importance of considering sex-shared and sex differences in gene expression. While we found some differences in the exact genes that show significant differences in gene expression, we observed that overall genes that showed sex differences in gene expression in term placentas also exhibited sex differences in late first trimester, and across most adult tissues, including all regions of the brain. Because of the comparisons we had the capacity to make here, we cannot fully distinguish the relative importance of sex chromosomes, sex hormones and gonadal sex on driving sex differences in gene expression in the samples studied here. Recent work in adult tissues (liver and adipose) in a mouse model reports the strongest effect of activational hormones, followed by a lesser organizational gonad effect, with a third strongest role of sex chromosome complement on sex differences in gene expression (Blencowe et al., 2022). Thus, one can hypothesize that a similar hierarchy may be in place for shaping sex differences in gene expression in humans. It is potentially provocative, then, that we observed that the most conserved sex differences across tissues and developmental timepoints were found on the X chromosome, suggesting a strong interaction between hormones, gonads, and chromosomes in determining sex differences in gene expression.

## Abbreviations

BMI: Body mass index
GATK: Genomic Analysis ToolKit
M: Million
MDS: Multidimensional scaling
SVD: Spontaneous vaginal delivery

## Declarations

### Ethics approval and consent to participate

This research was approved by HIC#1309012696 to The Yale University Reproductive Sciences Data and Specimen Biorepository, a component of the Department of Obstetrics, Gynecology & Reproductive Sciences, Yale School of Medicine, New Haven, CT.

### Consent for publication

Consent for publication was included in HIC#1309012696.

### Availability of data and materials

The datasets generated during this study are available via controlled access: phs002240.v1.p1 NIGMS Sex Differences Placentas at https://www.ncbi.nlm.nih.gov/projects/gap/cgi-bin/study.cgi?study_id=phs002240.v1.p1. The data from Gonzales et al was accessed here: NCBI GEO Accession GSE109082. The GTEx data was accessed here: https://storage.googleapis.com/gtex_analysis_v8/rna_seq_data/GTEx_Analysis_2017-06-05_v8_RNASeQCv1.1.9_gene_tpm.gct.gz. The code generated during this study is available at https://github.com/SexChrLab/Placenta_Sex_Diff.

## Funding

We acknowledge Research Computing at Arizona State University for providing high-performance computing and storage resources that have contributed to the research results reported within this paper (http://www.researchcomputing.asu.edu). We would also like to thank Wilson lab members for helpful feedback on the research and manuscript. This work was supported by the National Institute of General Medical Sciences (NIGMS) of the National Institutes of Health grant R35GM124827 to M.A.W. This research was supported by The Yale University Reproductive Sciences Data and Specimen Biorepository, HIC#1309012696, a component of the Department of Obstetrics, Gynecology & Reproductive Sciences, Yale School of Medicine, New Haven, CT. Research reported in this publication was supported by the Eunice Kennedy Shriver National Institute Of Child Health & Human Development of the National Institutes of Health under award number F31HD101252 to K.C.O. K.C.O. was additionally supported by ARCS Spetzler Scholar.

### Authors’ contributions

KCO: Conceptualization, Supervision, Formal Analysis, Investigation, Visualization, Writing - Original Draft, Writing - Review and Editing

SBP: Formal Analysis, Investigation, Visualization, Writing - Original Draft, Writing - Review and Editing

TNP: Conceptualization, Supervision, Visualization, Writing - Original Draft Preparation, Writing - Review and Editing

MS: Assistance with collecting the placenta samples

JO: Assistance with collecting the placenta samples

LP: Assistance with collecting the placenta samples

LR: Investigation

HJK: Conceptualization, Assistance with collecting the placenta samples, Writing – Review and Editing

MAW: Conceptualization, Supervision, Visualization, Resources, Project Administration, Writing - Original Draft Preparation, Writing - Review and Editing, Funding Acquisition

## Acknowledgements

We acknowledge Research Computing at Arizona State University for providing high-performance computing and storage resources that have contributed to the research results reported within this paper (http://www.researchcomputing.asu.edu). We would also like to thank Wilson lab members for helpful feedback on the research and manuscript.

