## Additional Table Legends and Figures for "Sex differences in early and term placenta are conserved in adult tissues"

### Additional Materials

#### Additional Tables Index

| | |
|---|---|
| Additional table 1 | Sample clinical information. |
| Additional Table 2 | Post-trimming sample sequence information. |
| Additional Table 3 | Samples removed from downstream analysis. |
| Additional Table 4 | Sex differences for clinical attributes. |
| Additional Table 5 | X and Y gametology gene list. |
| Additional Table 6 | Sex differences in innate immune genes. |
| Additional Table 7 | Sex differentially expressed genes. |
| Additional Table 8 | GTEx female and male mean TPM expression values. |
| Additional Table 9 | Gene FPKM and CPM values for late first trimester and term placentas. |
| Additional Table 10 | Unique and shared sex differentially expressed genes between first trimester and term placentas. |
| Additional Table 11 | Functional enrichment analysis for sex differentially expressed genes in term and first trimester placentas. |
| Additional Table 12 | Gene function annotation information and association with pregnancy complications for the first and term sex differentially expressed genes. |

#### Additional Table 1. Sample clinical information.

Clinical and sequence information for each full-term placenta sample.

#### Additional Table 2. Post-trimming sample sequence information.

Million sequences, percent of duplicate sequences, and percent QC content remaining after quality trimming.

#### Additional Table 3. Samples removed from downstream analysis.

Samples were removed that had less than 12.5M or higher than 90M sequences remaining after trimming. If more than 30% of the reads deviate from the sum of the deviations from the normal distribution of the per-sequence GC content as defined by the FASTQC report, then the sample was removed. If a sample clustered with opposite sex from the reported sex for that sample, then that sample was removed.

#### Additional Table 4. Sex differences for clinical attributes.

Sex differences for clinical information for full-term placentas for maternal age at delivery, pre-pregnancy BMI, gravidity and parity, gestational age, method of conception, self-reported race, and birth weight. Sex differences for continuous variables were tested using a t-test, p-value $< 0.05$. A Fisher's exact test was used to test for sex differences for categorical variables, p-value $< 0.05$.

#### Additional Table 5. X and Y gametology gene list.

A list of X and Y gametology genes were curated from a combination of Skaletsky et al. 2003 and Godfrey et al. 2020 (Godfrey et al., 2020; Skaletsky et al., 2003). FPKM expression for X-linked copy and Y-linked copy for all samples. In samples determined to have a Y chromosome, the FPKM value of the X-linked gametology and the Y-linked gametology were summed. expression between XX female X-linked gametology gene expression to XY male X-linked plus Y-linked gametology gene expression using a Wilcox rank-sum, p-value $\leq 0.05$.

#### Additional Table 6. Sex differences in innate immune genes.

979 innate immune games from InnateDB. Placenta CPM expression values for expressed innate immune genes in the late first trimester and term placentas. Sex differences in late first trimester and term placentas, adjusted p-value $\leq 0.05$.

##### Additional Table 7. Sex differentially expressed genes.

Sex differentially expressed genes in the late first trimester and term placentas, adjusted p-value < 0.05.

##### Additional Table 8. GTEx female and male mean TPM expression values.

Female and male mean TPM expression for 42 non-reproductive adult GTEx tissues. TPM expression for each gene obtained from counts version 2017-06-06_v8 (Carithers et al., 2015).

##### Additional Table 9. Gene FPKM and CPM values for late first trimester and term placentas.

Gene expression values for all genes provided in all included samples.

##### Additional Table 10. Unique and shared sex differentially expressed genes between first trimester and term placentas

Comparison of sex differentially expressed genes in term placentas and late first trimester placentas.

##### Additional Table 11. Functional enrichment analysis for sex differentially expressed genes in term and first trimester placentas.

Enriched biological processes, molecular functions, and gene families genes that had significantly higher expression in the female placenta samples or significantly higher expression in the male placenta samples.

##### Additional Table 12. Gene function annotation information and association with pregnancy complications for the first and term sex differentially expressed genes.

Sex differentially expressed genes annotated with molecular function and literature-search based association with pregnancy complications such as pre-eclampsia and miscarriage.

#### Additional Figure Index

| | |
|---|---|
| Additional Figure 1 | Sample sex check. |
| Additional Figure 2 | Multidimensional scaling plots reveal outlier samples. |
| Additional Figure 3 | Population ancestry inference. |
| Additional Figure 4 | Variation in expression trait attributes. |
| Additional Figure 5 | Sex differences for clinical attributes. |
| Additional Figure 6 | Sex differences in expression for gametolog genes. |
| Additional Figure 7 | Overlap of sex differentially expressed genes with and without birthweight as covariate. |

Additional Figure 1. Sample sex check.

Violin jitter plot CPM expression for each placenta sample for EIF1AY, KDM5D, UTY, DDX3Y, and RPS4Y1 Y-linked genes, and XIST, X-linked gene. Samples with at least two X chromosomes will show expression for XIST. Samples with the presence of the Y chromosome will show expression for Y-linked genes.

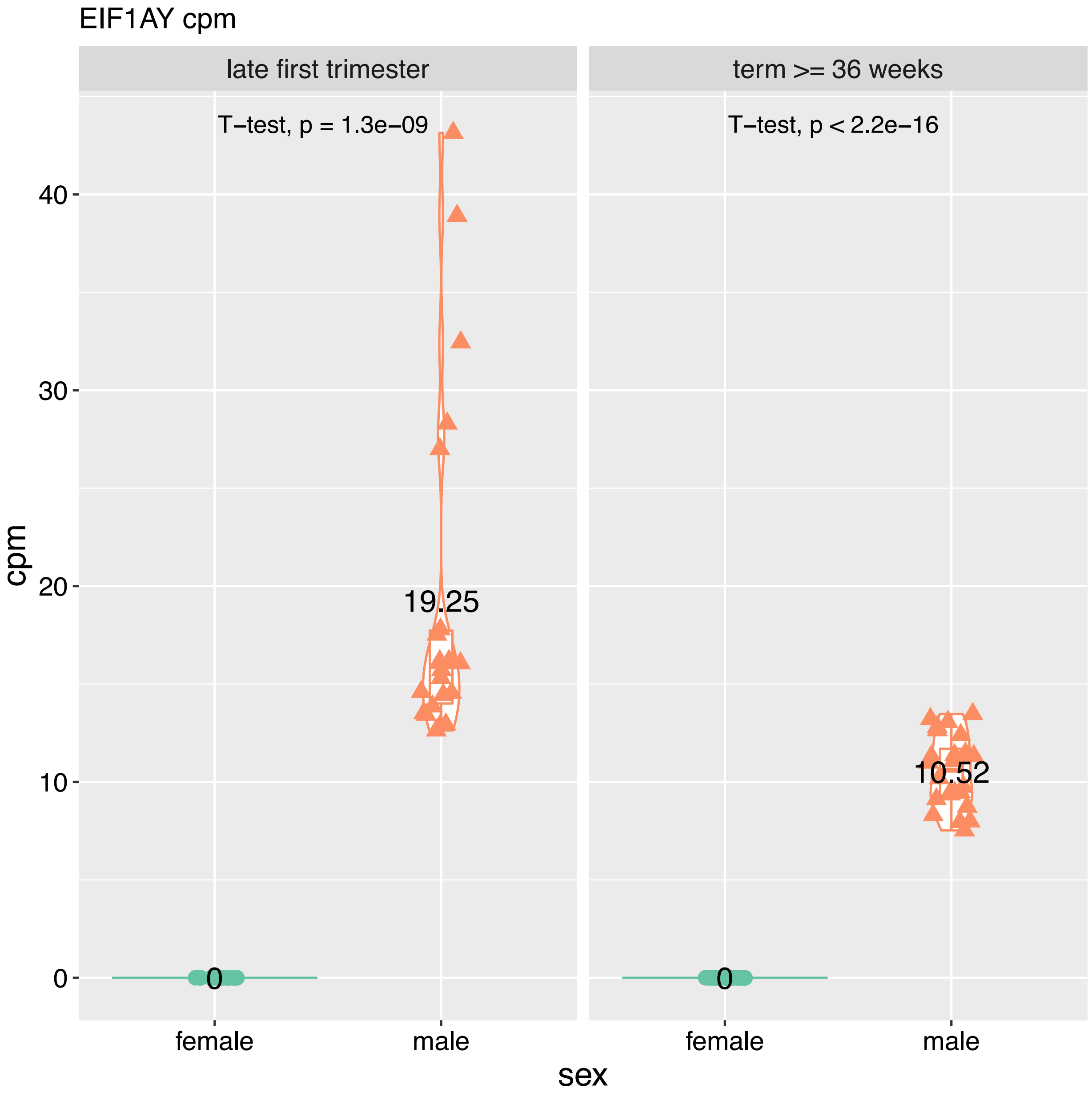

Additional Figure 2. Multidimensional scaling plots reveal outlier samples.

Multidimensional scaling (MDS) for all genes left and top too genes on the right for (A) late first trimester placentas (Gonzalez et al., 2018), (B) term placentas, (C) term placentas excluding failed samples.

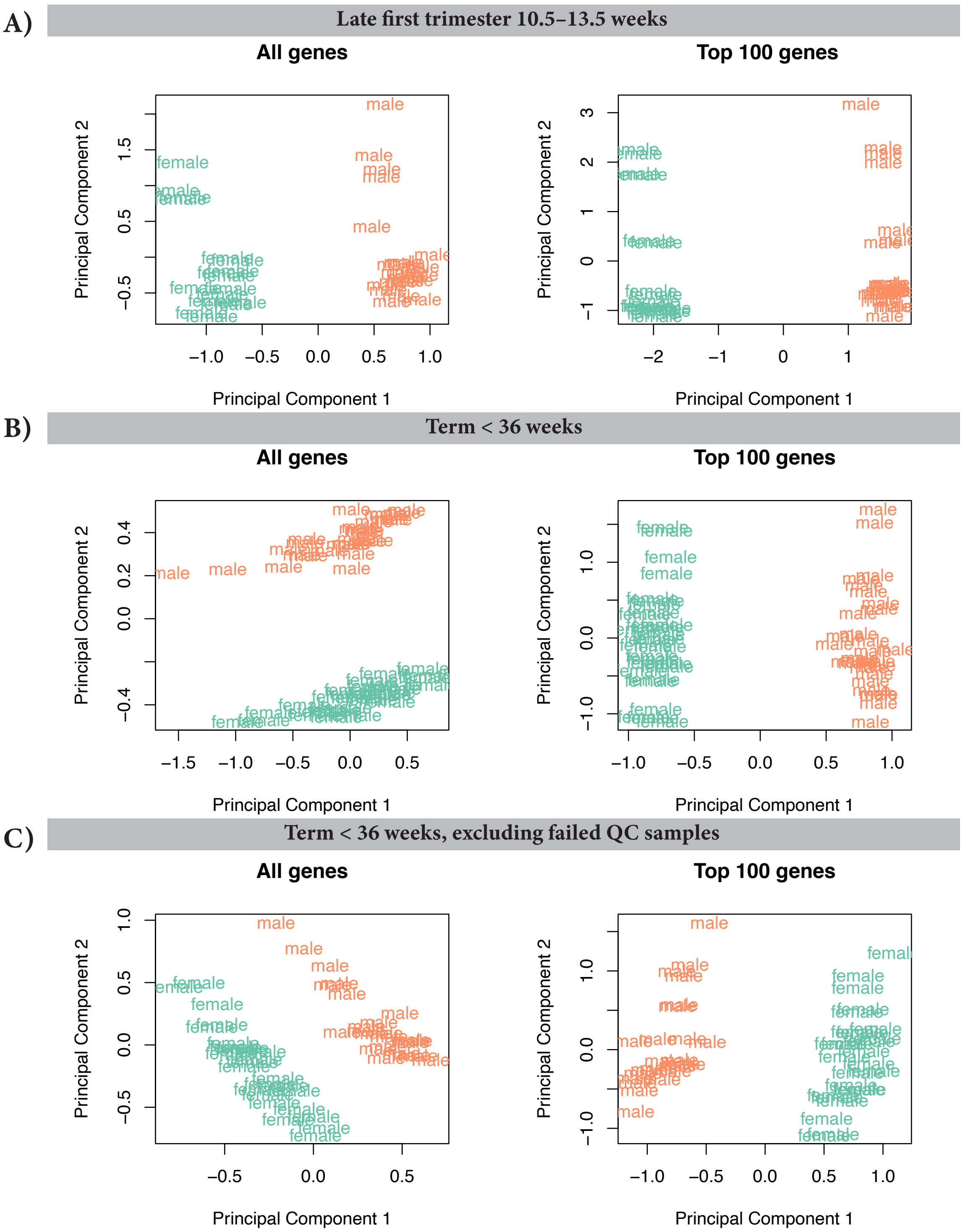

Additional Figure 3. Population ancestry inference.

Population ancestry was inferred from whole-exome sequencing for each full-term placenta.

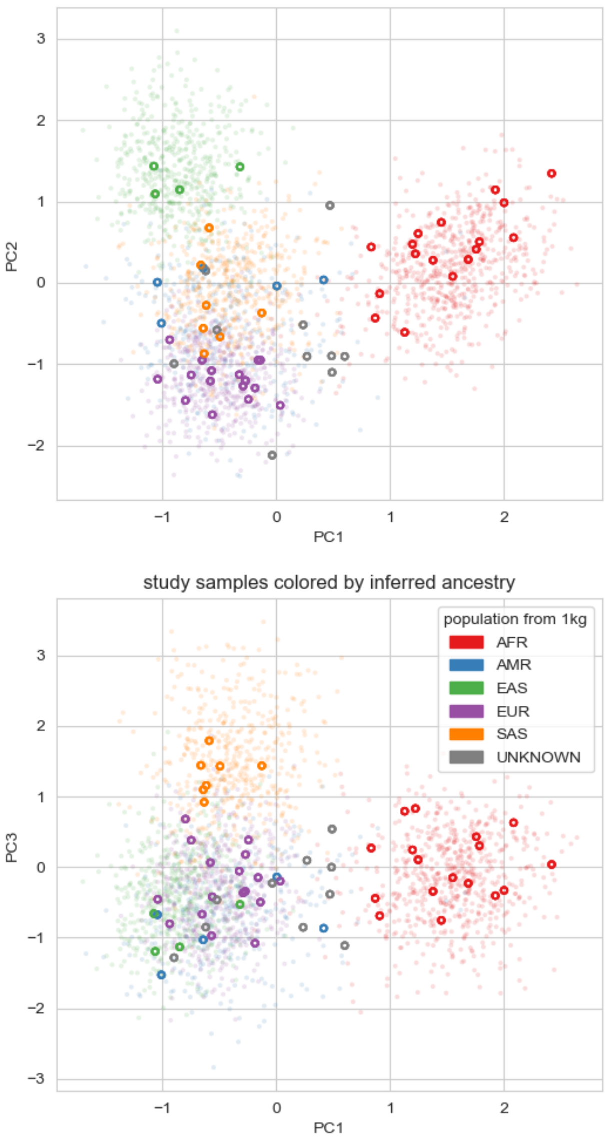

Additional Figure 4. Variation in expression trait attributes.

Variation within gestational age (GA), sequencing lane, sex, reported race, and birth weight was examined. Variation in placenta expression for maternal clinical data, including parity, gravidity, pre-pregnancy body mass index (BMI), and maternal age, were also examined.

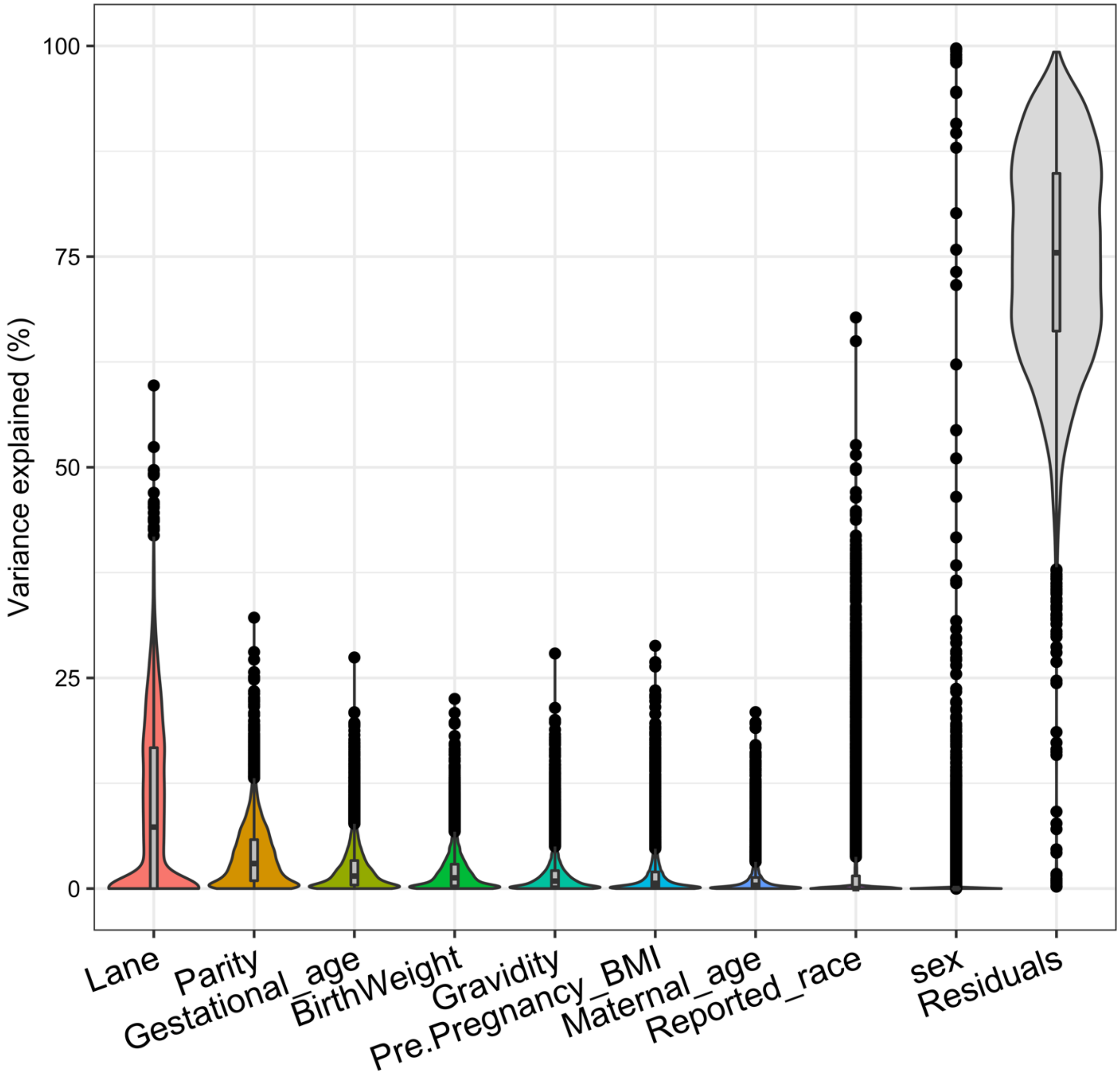

Additional Figure 5. Sex differences for clinical attributes.

Sex differences for clinical information for full-term placentas for maternal age at delivery, pre-pregnancy BMI, gravidity and parity, gestational age, method of conception, self-reported race, and birth weight. Sex differences for continuous variables were tested using a t-test, p-value < 0.05. A Fisher's exact test was used to test for sex differences for categorical variables, p-value < 0.05.

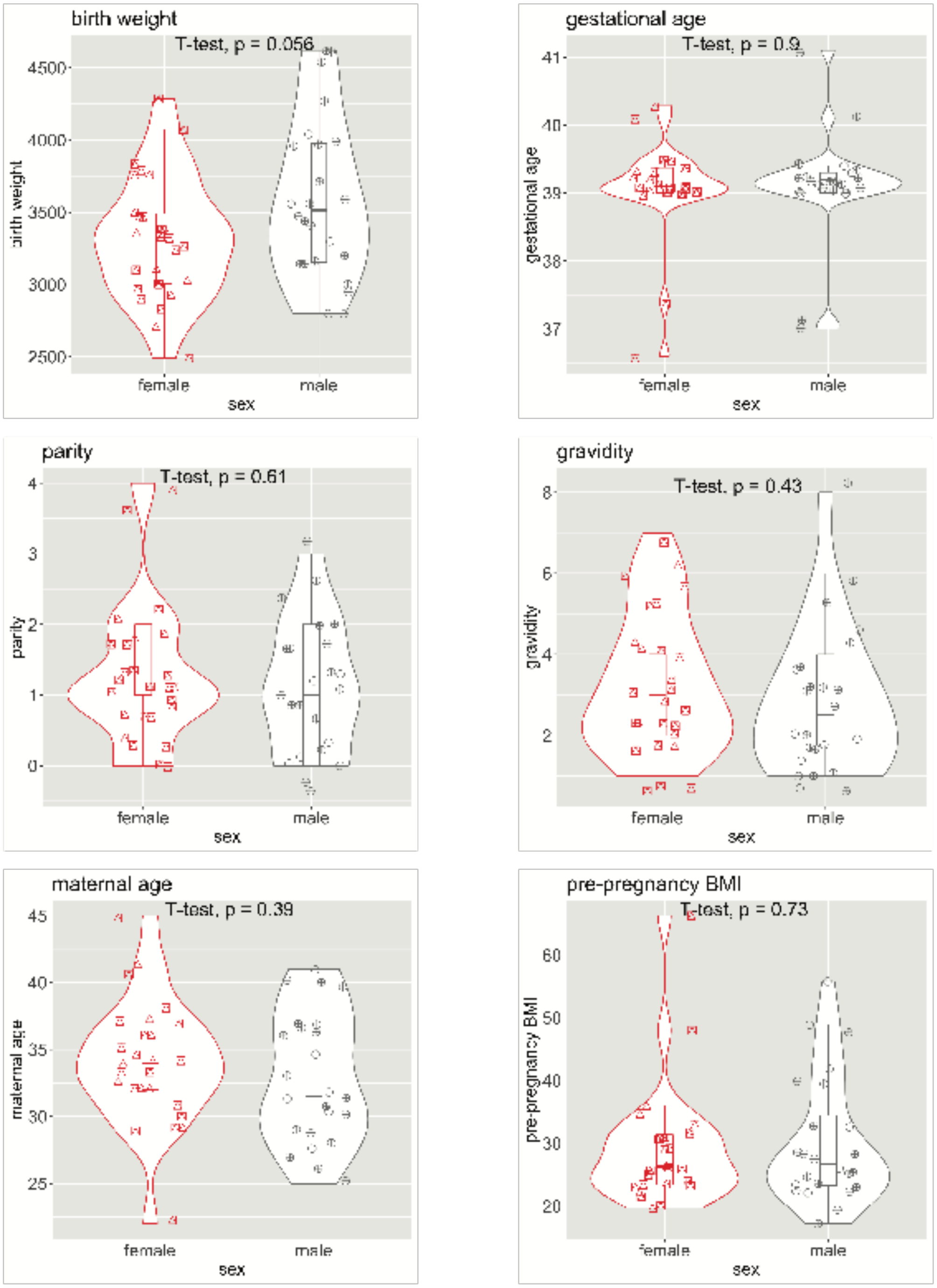

Additional Figure 6. Sex differences in expression for gametolog genes.

There is a significant difference in male XY to female XX expression for ZFX and KDM6A (UTX) when only looking at the X chromosome CPM expression value. When we add the Y chromosome-linked CPM expression count for these genes for male samples, there is no longer a difference in expression between males XY and females XX for ZFX. KDM6A, on the other hand, flips the measured sex-difference; it now shows males as having significantly higher expression than females. PCDH11X, when adding Y-linked CPM expression, shows a significantly higher expression than females. T-test to see if there is a difference between the female CPM and the male CPM for each gene, p-value < 0.05.

X+Y gene expression of gametolog genes in first trimester placentas:

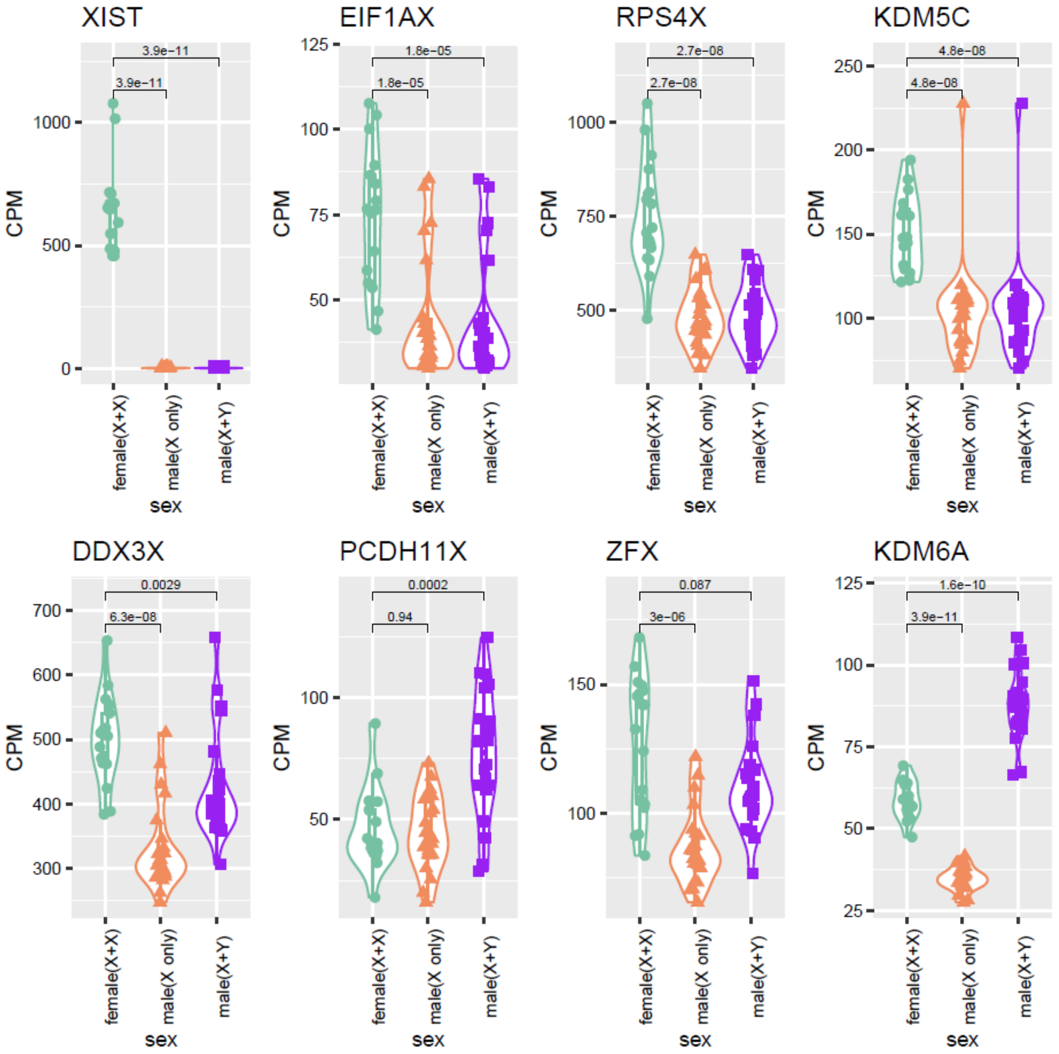

Additional Figure 7. Overlap of sex differentially expressed genes with and without birthweight as covariate.

Birthweight was added as a covariate in the linear model so that we can focus on gene expression changes based solely on genetic sex. 19 additional genes were identified as sex differentially expressed when differences in birthweight were not factored out; none had obvious connection to sex or growth and only 1 was consistently differentially expressed between first trimester and term placentas.

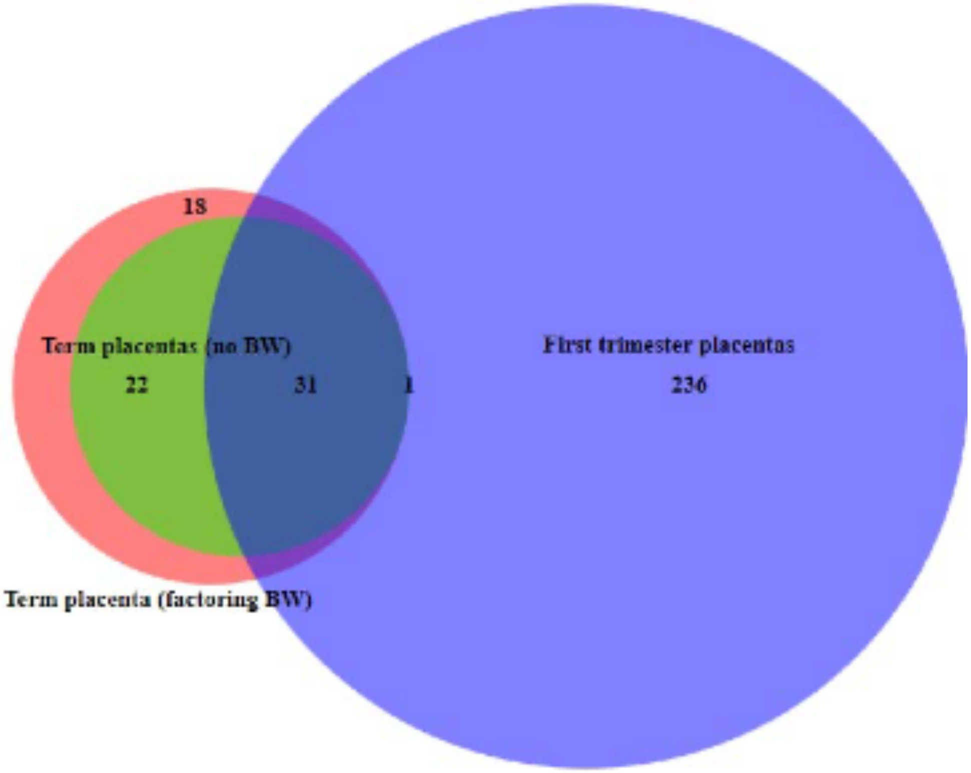
